# Therapeutic immunization against *Helicobacter pylori* infection in BALB/c mice induced by a multi-epitope vaccine based on computer-aided design

**DOI:** 10.1101/2021.02.28.433231

**Authors:** Junfei Ma, Shuying Wang, Qianyu Ji, Qing Liu

## Abstract

**Background:** Combined antibiotic regimens have caused problems such as increasing antimicrobial resistance to *H. pylori* and intestinal flora disturbance. Vaccination is a great alternative approach, but also faces the limited immune response induced by monovalent vaccines. Therefore, the development of multi-epitope vaccines is promising immunotherapy to control *H. pylori* infection.

**Objective:** To develop a multi-epitope vaccine and evaluate its therapeutic efficacy against *H. pylori* infection.

**Materials and Methods:** The B and T cell epitopes from UreB, FlaA, AlpB, SabA, and HpaA were linked for producing 2 multi-epitope vaccines (CTB-S3 and CTB-S5) by a structural evaluation based on computer-aided design. The abilities to produce antigen-specific antibodies and neutralizing antibodies of CTB-S3 and CTB-S5 were evaluated in BALB/c mice. After that, their therapeutic efficacy was explored in *H. pylori-infected* mice.

**Results:** CTB-S3 or CTB-S5 could induce high levels of specific antibodies against UreB, FlaA, AlpB, SabA, HpaA, and neutralizing antibodies against *H. pylori* urease and adhesion. Also, oral therapeutic immunization with CTB-S3 or CTB-S5 could decrease *H. pylori* colonization and reduce stomach damage; the protection was correlated with *H. pylori*-specific IgG, SIgA antibodies, and CD4^+^ T cell immune response.

**Conclusions:** Our study developed a multi-epitope vaccine based on a computer-aided design. The CTB-S3 and CTB-S5 vaccines may be promising therapeutic candidate vaccines against *H. pylori* infection and provide a reference for vaccine design of other pathogens.

## 1| INTRODUCTION

*Helicobacter pylori* (*H. pylori*) is the most important etiologic factor for gastric diseases including chronic gastritis, peptic ulcers, and gastric malignancies^1,2^. To date, the major antibiotic-based triple regimen has caused the increased antimicrobial resistance and intestinal flora disturbance^3,4^. Therefore, the development of potential vaccines is a tremendous priority for controlling *H. pylori* infection.

A range of antigens has been developed to construct the vaccine candidates^5^. Urease is a key functional protein that could neutralize stomach acidity and promote chemotaxis by decomposing urea, which helps *H. pylori* to colonize the stomach^6,7^. The strongly immunogenic subunit UreB is a widely used antigen, whose B cell epitopes and CD4^+^ T cell epitopes were identified in the previous studies^8–12^. The movement-related protein flagellin A (FlaA) is also a candidate antigen, whose B cell epitope was identified^13^. Besides, adhesins play a critical role in *H. pylori* attachment to the gastric mucosa; so sialic acid-binding adhesin (SabA), hop family adhesin AlpB (AlpB), and *H. pylori* adhesin A (HpaA) have been identified as excellent candidate antigens to develop *H. pylori* vaccines^14^. Their possible conservative adhesion domains^15–17^ are crucial for the production of neutralizing antibodies against *H. pylori* adhesion to the gastric mucosa.

Due to the limited immune response induced by monovalent vaccines, the development of multi-epitope vaccines is becoming the focus of immunotherapy against *H. pylori*. Current studies^18,19^ have shown that the multi-epitope vaccines could generate a broadly reactive antibody response against *H. pylori* antigens and significantly reduced *H. pylori* colonization in the stomach, indicating that the multi-epitope vaccine is a promising vaccine strategy. In the construction of a multi-epitope vaccine, the B and T cell epitopes with different permutations could produce different constructs. These different constructs make a difference in the stability of the recombinant vaccine and the exposure of the critical domains. The efficient immune response induced by the proper antigen is strongly dependent on the optimal structural stability of the vaccine protein^20^. Therefore, the assembly and permutation of epitopes are the keys to the successful design of the multi-epitope vaccine.

Computer-aided design could invoke available structure information to construct immunogens based on immunoinformatics^21^. By the evaluation of the predicted structure and other physicochemical features including hydrophilicity-hydrophobicity, solubility, and pI, the construct with optimal epitope permutation will be selected for further experimental verification^22^. In addition, some recent studies^23–25^ have reported the design of multi-epitope vaccines based on computer aid. However, these studies are still at the design stage, which means the effectiveness of the computer-aid multi-epitope vaccines against *H. pylori* has not been verified.

In this study, we constructed 2 multi-epitope vaccines (CTB-S3 and CTB-S5) with epitopes from UreB, FlaA, AlpB, SabA, and HpaA based on computer-aided design especially structural evaluation. The oral CTB-S3 or CTB-S5 vaccines could induce broad specific antibody response, reduce *H. pylori* loads, and protect the stomach. These findings suggest that the multi-epitope vaccine designed by computer aid represents a promising approach for the development of an effective *H. pylori* vaccine and provides a novel strategy for the design of multi-epitope vaccines against other pathogens.

## 2| Materials and methods

### 2.1| Plasmids, bacteria, and animals

The plasmid pEC was a gift from Dianfan Li of the National Center for Protein Science. Compenent cells *Escherichia coli* (*E. coli*) DH5α and *E.coli* BL21(DE3) were purchased from Vazyme (Nanjing, China) and cultured in LB medium.

The mouse-adapted *H. pylori* strain SS1 was obtained from Shanghai Institute of Digestive Disease and preserved in our laboratory. *H. pylori* was cultured on Columbia agar plates enriched with 7% new-born calf serum, polymyxin B (5 μg/mL), trimethoprim (5 μg/mL), and vancomycin(10 μg/mL) under microaerophilic conditions (5% O_2_, 10% CO_2_, and 85% N_2_) at 37 °C for 3-5 days.

Specific pathogen-free (SPF) BALB/c mice, 5–6 weeks of age, 14±2 g, were purchased from Jiesijie (Shanghai, China). The mice were allowed 1 week to adapt to the environment before starting the experiments. This study was approved by China Ethics Committee.

### 2.2| Design of the multi-epitope vaccines

The selected epitopes were listed in Table 1. Among them, the epitopes from UreB and FlaA were obtained from the Immune Epitope Database (IEDB) and the B cell epitopes from *H. pylori* adhesins (HpaA, AlpB, and SabA) were predicted by BepiPred 2.0 Server^26^. Particularly, three epitopes from adhesins contain the previously reported adhesion domains^15–17^.

**TABLE 1.**
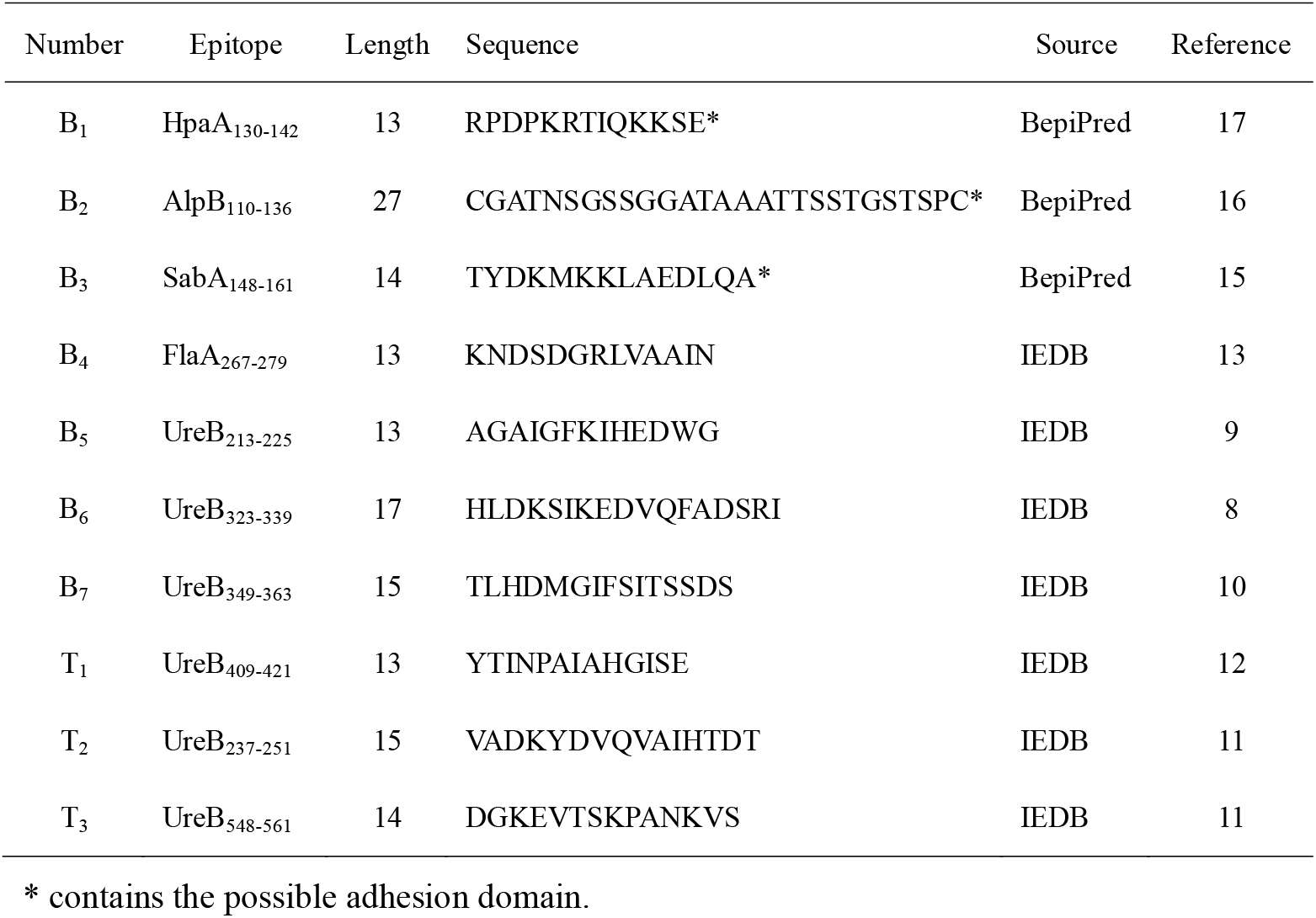
The epitopes from UreB, FlaA, HpaA, AlpB, and SabA for the multi-epitope vaccine construction.

The link of the same epitopes in different permutations could produce different constructs. Twenty constructs of tandem copies including 7 B cell epitopes, 3 T cell epitopes, and the linkers “KK”, “GS” were determined randomly. The selected linkers could effectively connect the epitopes without changing the immunogenicity of designed epitopes^27,28^. After different attempts based on elementary evaluation of VaxiJen 2.0^29^(VaxiJen score > 1.10), 7 constructs with higher VaxiJen scores numbered S1-S7 were obtained. Subsequently, the sequences of 7 constructs were submitted to the I-TASSER sever^30^ to predict their 3D structures. Their sequences were listed in Table S1.

### 2.3| Structure validation of the multi-epitope vaccine constructs

ProSA-web and RAMPAGE sever were used for evaluating 7 multi-epitope vaccine constructs based on their predicted structures. The Z-score calculated by ProSA-web could assess the overall quality of the predicted structure^31^. The main Ramachandran plot from RAMPAGE was used for calculating phi–psi torsion angles for each amino acid in the vaccine structure^32^.

### 2.4| Construction of multi-epitope vaccines for oral immunization

To enhance the mucosal immunogenicity of screened multi-epitope vaccines, cholera toxin subunit B (CTB), a widely used mucosal adjuvant, was added to the N-terminus of the screened Construct S3 or S5 with a flexible linker “DPRVPSS”. After amplification, cloning, and transformation, CTB-S3 and CTB-S5 were expressed by *E. coli* BL21 (DE3). The proteins were purified by affinity chromatography and assessed by SDS-PAGE. Their immunoreactivity was evaluated by western blotting.

CTB could bind intestinal epithelial cells and antigen presenting cells through monosialotetrahexosylganglioside (GM1) receptors, which then mediates antigen entry into the cell^33^. GM1-ELISA was used to demonstrate the adjuvanticity of CTB component in CTB-S3 or CTB-S5 as previously described^34^. Briefly, ELISA plates were coated with GM1 ganglioside at 4 □ for 12 h. After washing, ELISA plates were blocked by incubating with 5% (m/V) skim milk for 2 h. The CTB-S3, CTB-S5, or CTB protein was added to the plates and incubated at 37 □ for 2 h. After washing, a proper dilution (1:1000) of anti-CTB mouse monoclonal antibody (Sigma, USA) was added to the plates and incubated at 37 □ for 1 h. After washing, HRP-conjugated goat anti-mouse IgG (Invitrogen, USA) was added to the plates and incubated at 37 □ for 1 h. Substrate tetramethylbenzidine (TMB) was then added and incubated for 15 min. The absorbance was measured at 450 nm.

### 2.5| Subcutaneous immunization

SPF BALB/c mice were randomly divided into 3 groups (n=5) and were vaccinated subcutaneously 3 times at 7-day intervals with 100 μg of the purified CTB, CTB-S3, or CTB-S5 in complete Freund’s adjuvant (FA, Sigma, USA) on day 0 and in incomplete FA on the 7th and 14th day. The pure proteins CTB, CTB-S3, or CTB-S5 were used as immunogens in the last booster immunization on the 21st day. Serum was collected at 7 days after the last vaccination to determine the specific antibodies.

### 2.6| Determination of specific antibodies after subcutaneous immunization

ELISA plates were coated with 1 μg/well of each antigen (UreB, FlaA, SabA, AlpB, or HpaA) respectively at 4 □ overnight. After washing, the plates were blocked with 5% (m/V) skim milk for 2 h. The diluted serum samples (1:1000) were added to the antigen-coated plates and incubated for 1 h. After washing, a proper dilution of HRP-conjugated goat anti-mouse IgG was added to the plate and incubated for 1 h. Finally, TMB was added and incubated at room temperature for 15 min. The reaction was then terminated with 2 M H_2_SO_4_. The absorbance was measured at 450 nm using a microplate reader.

### 2.7| Assay of neutralizing antibodies against *H. pylori* urease and adhesins

Serum IgG antibodies were purified by protein G column chromatography (GE Healthcare, USA). The purified IgG antibodies were detected by SDS-PAGE. To determine neutralizing antibodies against *H. pylori* urease, *H. pylori* urease (2 μg in 50 μL, Creative Enzymes, USA) was incubated with purified IgG antibodies (64 μg/well) in 96-well microtiter plates overnight at 4 °C. After that, 100 μL of 50 mM PBS containing 500 mM urea, 0.02% phenol red, and 0.1 mM dithiothreitol (DTT) was added to each well. The absorbance was measured at 550 nm. Percentage inhibition of urease activity = [(activity without antibodies – activity with antibodies)/(activity without antibodies)] × 100 %.

To determine the neutralizing antibodies against *H. pylori* adhesins, the AGS human gastric cancer cell adhesion assay was carried out on *H. pylori* SS1 using a CFU counting method. Approximate 2×10^5^ AGS cells were seeded in a 12-well plate per well overnight. Approximate 2×10^7^ CFU *H. pylori* cultures were incubated with 40 μL of purified IgG antibodies (100 μg/mL) and slightly shaken at room temperature for 2 h. AGS cells were washed three times with PBS, and the *H. pylori* incubates were added to the wells at a multiplicity of infection of 100. The mixtures were incubated in a CO_2_ incubator at 37 °C for 2 h. After incubation, the wells were washed 3 times with PBS containing 1% saponin. After the mechanical treatment, the mixtures were plated onto Columbia agar for bacteria counting. Percentage inhibition of adhesion = [(CFU without antibodies – CFU with antibodies)/(CFU without antibodies)] × 100 %.

### 2.8| Infection and therapeutic vaccination

SPF BALB/c mice (male, 5-6 weeks old) were infected with *H. pylori* SS1(10^9^ CFU/mouse) intragastrically, four times within two weeks. Four weeks after the last infection, the *H. pylori*-infected mice were randomly divided into 3 groups (n=5) and vaccinated intragastrically with 100 μg of antigen (CTB, CTB-S3, or CTB-S5) in 0.2 M sodium hydrogen carbonate buffer (200 μL) for four times at 1-week interval. Two weeks after the final immunization, the mice were sacrificed and examined. The whole therapeutic vaccination procedure was showed in Figure 1.

**FIGURE 1.**
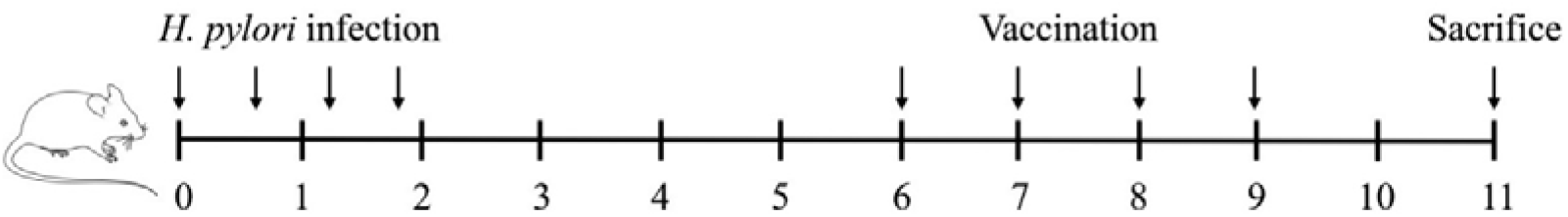
The procedure of *H. pylori* infection and therapeutic vaccination.

### 2.9| Detection of specific IgG in serum and SIgA in the stomach mucosa

The specific IgG in serum and secretory IgA (SIgA) in stomach mucosa against *H. pylori* were determined by ELISA. ELISA plates were coated with 5μg/mL *H. pylori* lysates at 4 □ overnight. To determine specific IgG, the antisera were collected and diluted 1:1,000 in PBS. To determine stomach SIgA, one-fourth of stomach tissue was homogenized in 1 mL of PBS containing 0.1 mM Phenylmethanesulfonyl fluoride (PMSF). The supernatant was collected and diluted 1:5 in PBS. An HRP-conjugated goat anti-mouse IgG or an HRP-conjugated goat anti-mouse IgA (Sigma, USA) was used as the secondary antibody.

### 2.10| Examination of *H. pylori* colonization in stomachs

To examine the *H. pylori* colonization in stomachs, one-fourth of stomach tissue was weighed and homogenized in 1 mL of PBS. Serial 10-fold dilutions of the stomach homogenate were plated on Columbia agar supplemented with 7 % new-born calf serum and *H. pylori* selective supplement (Oxoid, UK). After cultured for 3-5 days at 37 □, colonies were counted and the number of CFU per stomach was calculated.

### 2.11| Gastric histology

One fragment of stomach tissue was fixed with formalin, embedded in paraffin, and stained with hematoxylin and eosin (HE) according to the standard procedure^35^.

### 2.12| Cytokine production

To determine cytokine production, the splenic lymphocytes were isolated and cultured (2×10^5^ cells/well) with *H. pylori* lysates (5 μg/mL) in 12-well plates at 37 □ for 72 h. The culture supernatants were collected for the determination of IFN-γ, IL-4, and IL-17 using ELISA kits (Jiang Lai Biotech, Shanghai, China) according to the manufacturer’s instructions.

### 2.13| Statistical analyses

All independent experiments carried out in this study and showed in figures were biological replicates. All data were analyzed with GraphPad Prism software using One-way ANOVA. P < 0.05 was considered statistically significant. **p < 0.01, *** p < 0.001, ns, not significant).

## 3| RESULTS

### 3.1| Design and screening of the multi-epitope vaccines

Seven constructs were preliminarily screened based on the VaxiJen score (Figure S1 A). Further, their structures were predicted by I-TASSER server (Figure S1 B). To evaluate the structural rationality of 7 constructs, ProSA-web and RAMPAGE sever were used for the structure validation. As shown in Table 2, the C-score of Construct S3 or S5 was the top 2 of all the constructs. Z-score of Construct S3 and S5 were −2.39 and −0.96, which is in the range of native protein conformation scores (Figure 2 B and E). Ramachandran plot from RAMPAGE showed that the proportion of the amino acids in the favored and allowed regions of Construct S3 and S5 were 91.7% and 94.1%, which performed well in all the constructs (Figure 2 C and F). All these results indicated that the structures of Construct S3 and S5 were more stable and reasonable. The structures of Construct S3 and S5 were shown in Figure 2 A and D.

**FIGURE 2.**
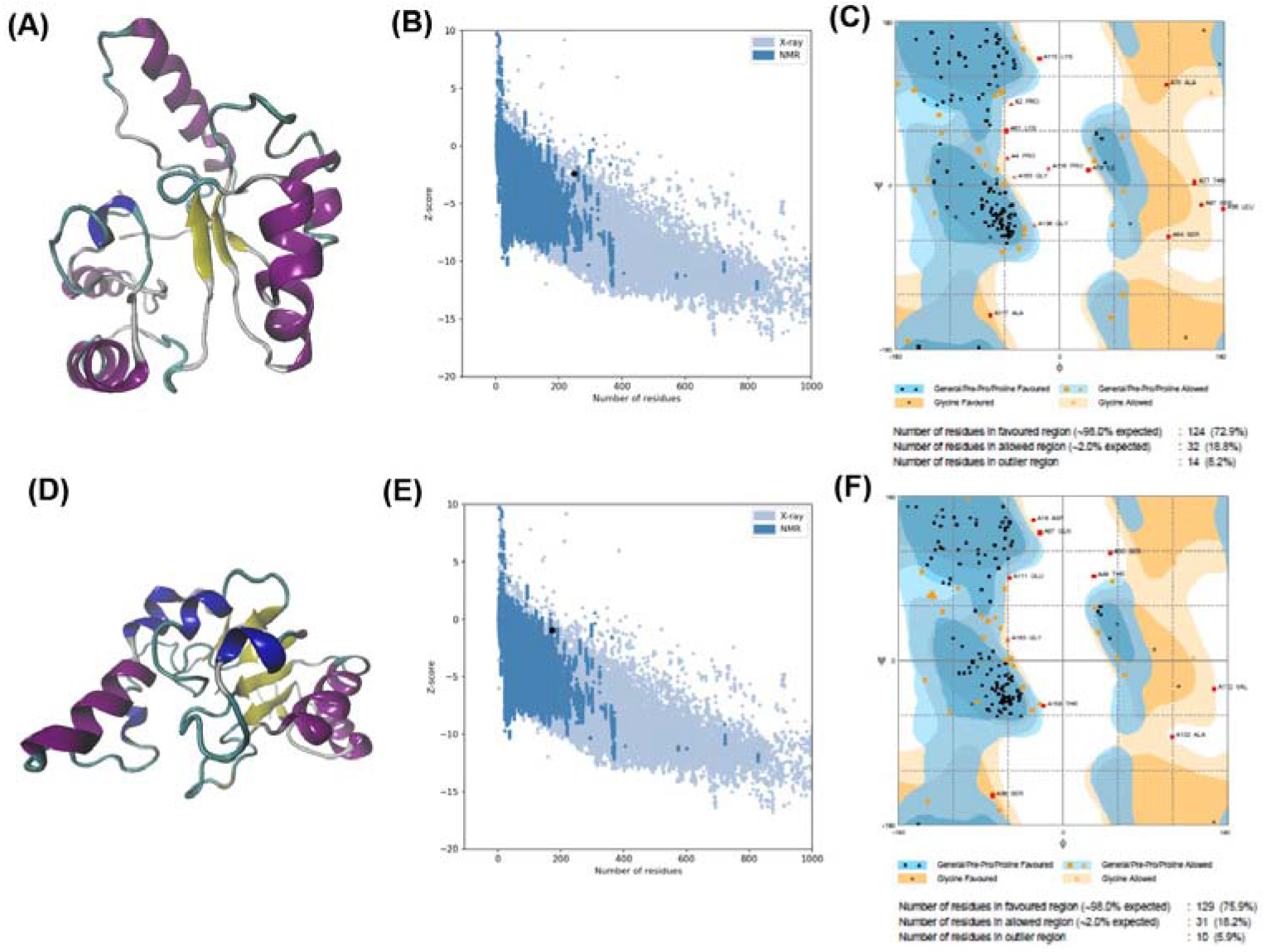
Structural prediction and evaluation of the screened Construct S3 and S5. (A) The predicted structure of Construct S3 by I-TASSER. (B) The z-score plot of the predicted S3 structure by ProSA-web. (C) Ramachandran plot analysis of the predicted S3 structure. (D) The predicted structure of Construct S5 by I-TASSER. (E) The z-score plot of the predicted S5 structure by ProSA-web. (F) Ramachandran plot analysis of the predicted S5 structure.

**TABLE 2.**
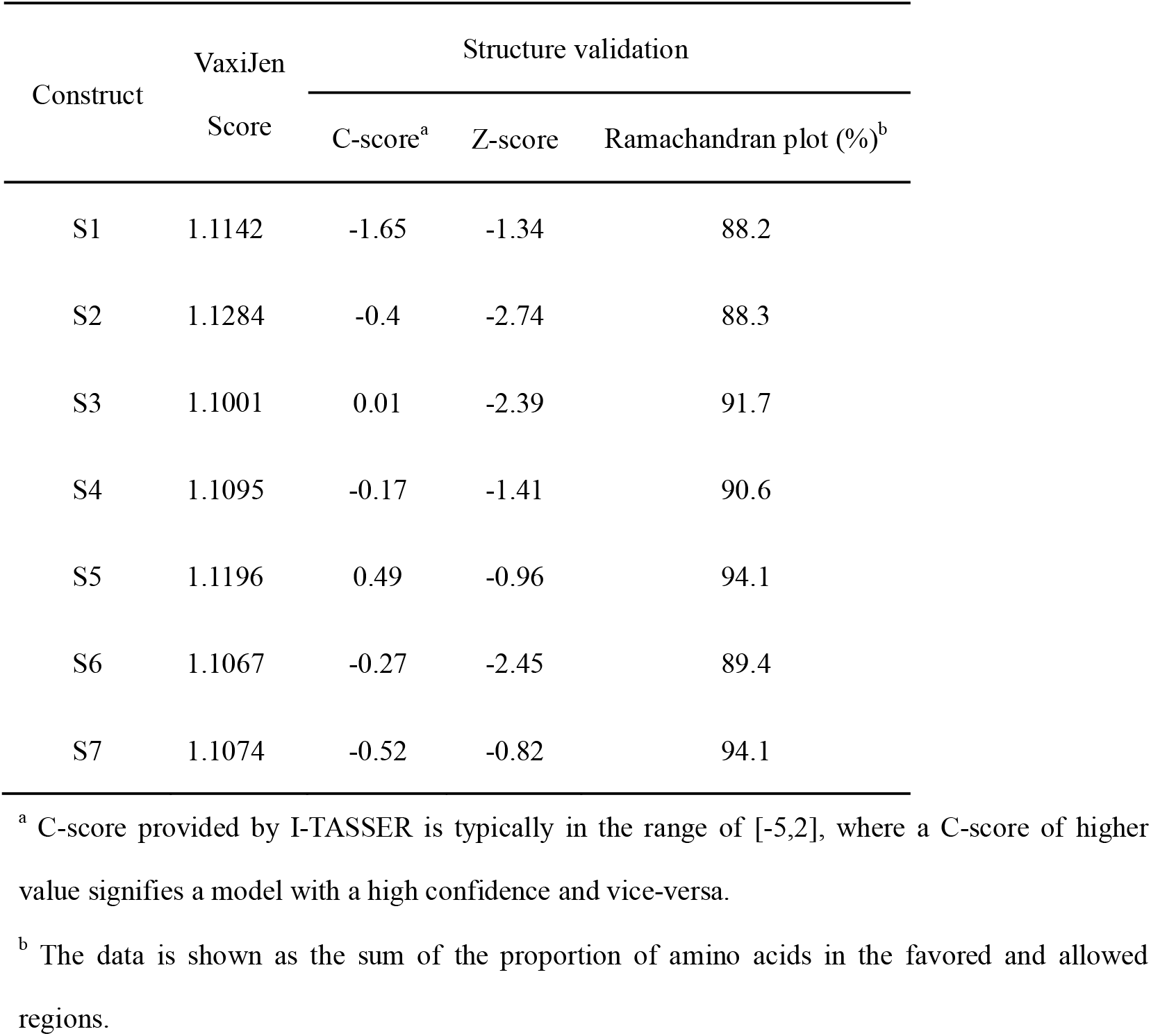
Structure validation of the 7 constructs.

### 3.2| Construction of multi-epitope vaccines for oral immunization

According to the schematic representation (Figure 3 A), CTB-S3 or CTB-S5 immunogen was constructed by the addition of CTB to the N-terminal of S3 or S5 with a flexible linker “DPRVPSS” The CTB-S3 and CTB-S5 proteins were successfully expressed and purified (Figure 3 B). They could be identified by anti-His mouse polyclonal antibody (Figure 3 C). The purified CTB-S3 and CTB-S5 proteins both display 2 bands. Since the His-tag is located at the C-terminal of the proteins, it may be caused by the N-terminal degradation of part of the proteins. Further, the adjuvanticity of CTB in CTB-S3 or CTB-S5 was analyzed by GM1-ELISA. CTB-S3 and CTB-S5 proteins were able to bind the coating GM1 as CTB (Figure 3 D), which indicated that CTB in CTB-S3 and CTB-S5 proteins could play the adjuvanticity.

**FIGURE 3.**
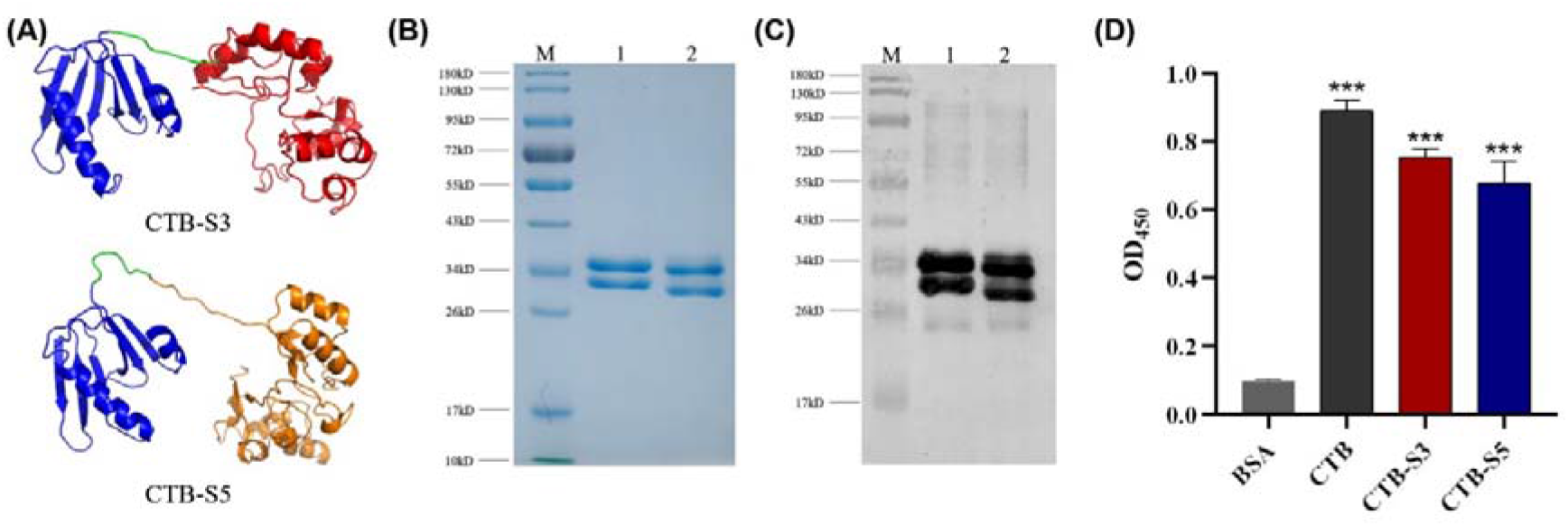
Construction of CTB-S3/S5 for oral immunization. (A) The construction diagram of CTB-S3/S5. CTB (Blue) and S3 (Red) or S5 (Orange) were linked with a linker “DPRVPSS” (Green). (B) Expression and purification of CTB-S3/S5 proteins visualized by SDS-PAGE. M: Marker; 1, CTB-S3; 2, CTB-S5. (C) Expression and purification of CTB-S3/S5 proteins visualized by Western blotting. CTB-S3/S5 proteins could be recognized by anti-His mouse monoclonal antibody. M: Marker; 1, CTB-S3; 2, CTB-S5. (D) The adjuvant effect of CTB in CTB-S3/S5 by GM1-ELISA. These results were verified in triplicate assays. ***p < 0.001.

### 3.3 Determination of antigen-specific antibodies and neutralizing antibodies after subcutaneous immunization

After subcutaneous immunization with CTB, CTB-S3, or CTB-S5, the antigen-specific antibodies in the antiserum were assayed by ELISA. Compared with CTB, CTB-S3 or CTB-S5 could induce a high level of antibodies against antigens including UreB, FlaA, AlpB, SabA, and HpaA (Figure 4 A). And there was no significance between CTB-S3 and CTB-S5 groups at the level of antigen-specific antibodies. It indicated that the B cell epitopes in CTB-S3 or CTB-S5 construct have good immunogenicity.

**FIGURE 4.**
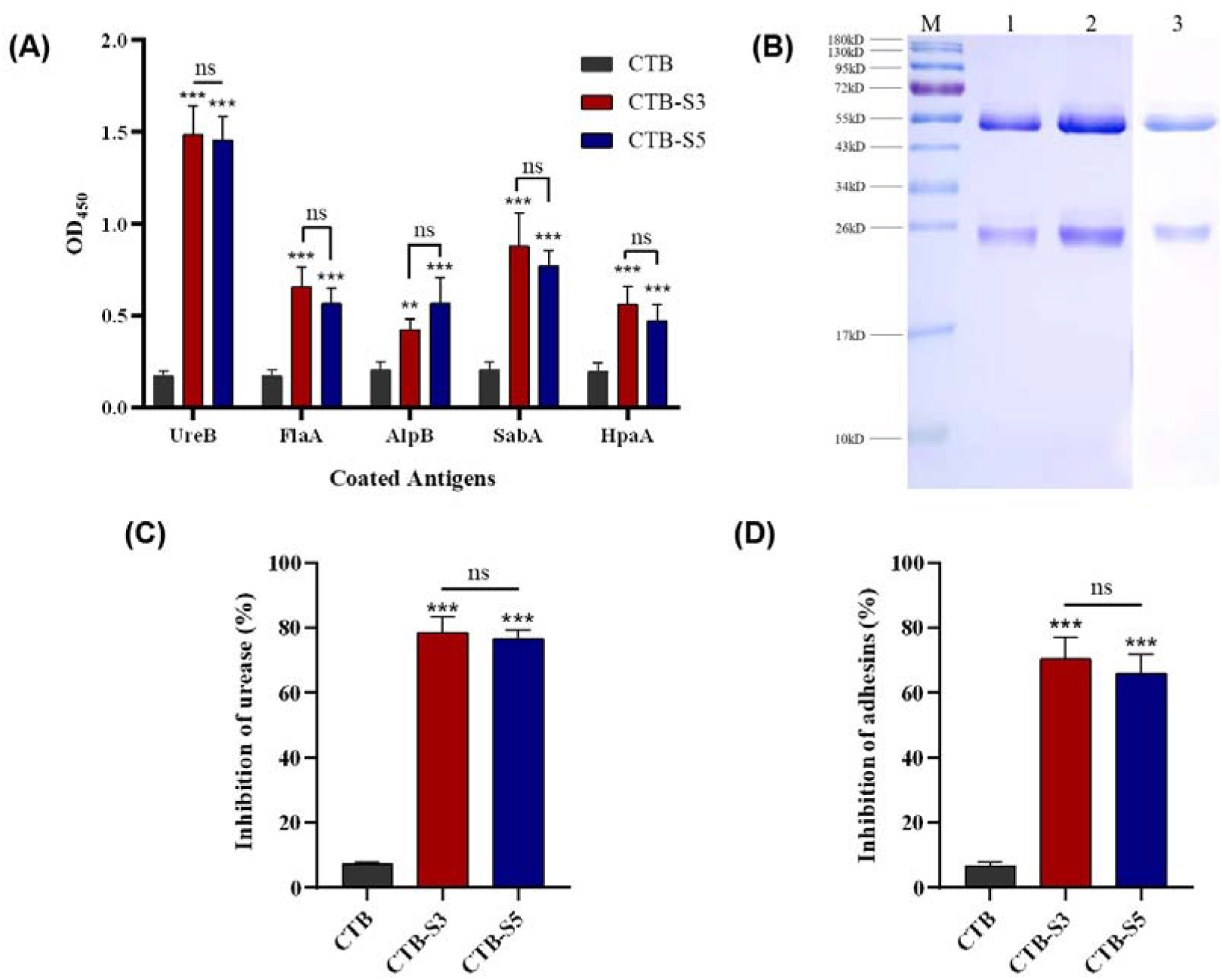
Evaluation of antibody response generated by CTB, CTB-S3, or CTB-S5 after subcutaneous immunization (n=5). (A) Determination of antigen-specific antibodies in serum by ELISA after subcutaneous immunization with CTB, CTB-S3, or CTB-S5. (B) Visualization of IgG antibodies purified from the serum of mice immunized by CTB, CTB-S3, or CTB-S5. M, Marker; 1, CTB-S3; 2, CTB-S5; 3, CTB (C) *H. pylori* urease neutralization test of the purified IgG. (D) Adherence inhibition assay of the purified IgG. Data are mean ± S.D. ***p < 0.001; **p < 0.01; ns, not significant.

To evaluate the neutralizing antibodies against *H. pylori* urease and adhesins, mouse IgG antibodies in antiserum were purified and then identified by SDS-PAGE (Figure 4 B). Compared with anti-CTB IgG, anti-CTB-S3 or anti-CTB-S5 IgG could significantly inhibit the enzyme activity of *H. pylori* urease (Figure 4 C) and *H. pylori* adhesins to AGS cells (Figure 4 D). Also, there was no significance between anti-CTB-S3 and anti-CTB-S5 IgG in the inhibitions of *H. pylori* urease and adhesins.

### 3.4| Evaluation of therapeutic vaccination

After oral therapeutic vaccination, the specific IgG and SIgA antibodies against *H. pylori* were analyzed by ELISA. Compared with CTB, CTB-S3 or CTB-S5 could induce a higher level of *H. pylori*-specific IgG antibody in serum (Figure 5 A) and SIgA antibody in the stomach (Figure 5 B). In addition, there was no significance between CTB-S3 and CTB-S5 at the levels of serum IgG and stomach SIgA antibodies. Further, the *H. pylori* colonization in the stomach was analyzed by quantitative culture. Compared with CTB, the immunization with CTB-S3 or CTB-S5 significantly decreased the *H. pylori* loads in the stomach (Figure 5 C). And there was no significance between CTB-S3 and CTB-S5 in *H. pylori* colonization.

**FIGURE 5.**
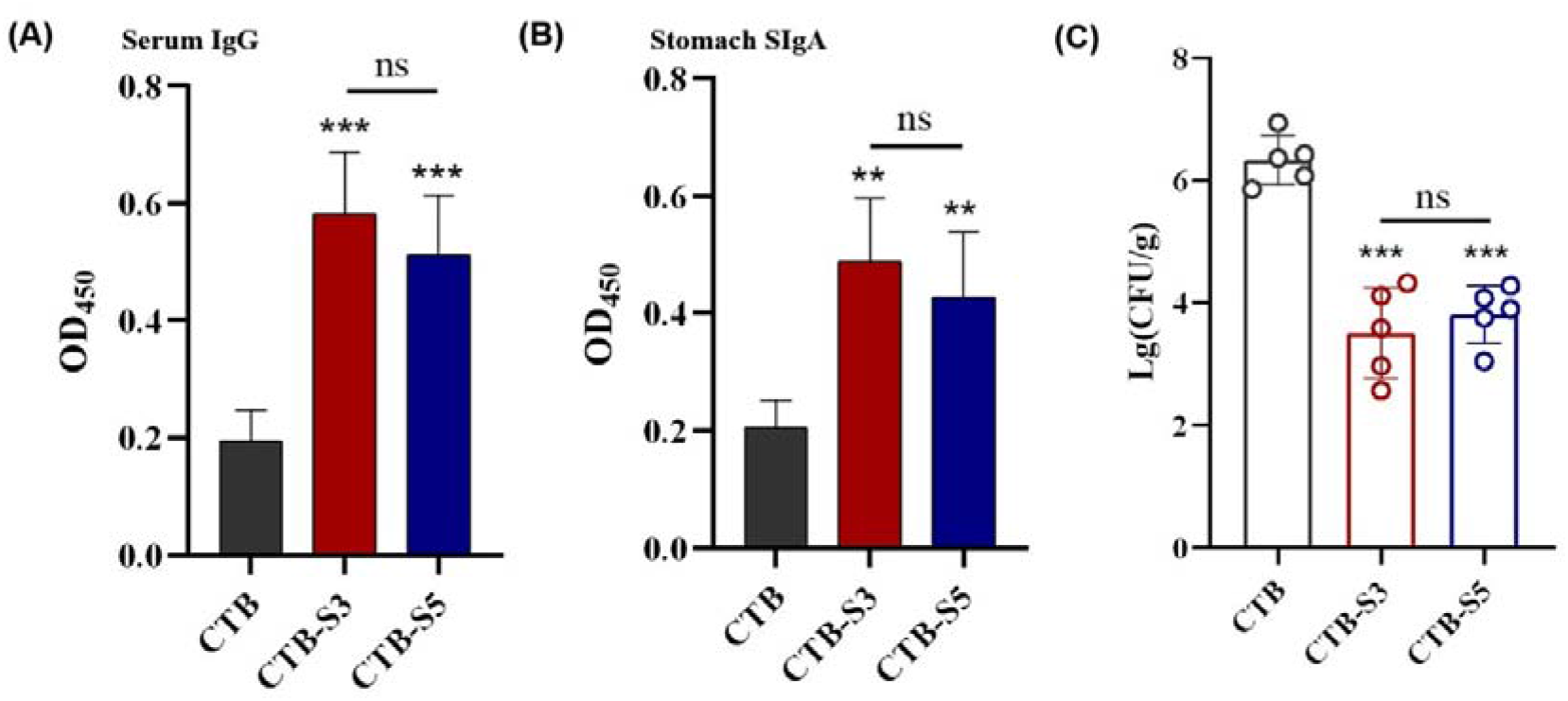
Evaluation of therapeutic vaccination with CTB, CTB-S3, or CTB-S5. (B) Determination of serum IgG antibody against *H. pylori* lysates after therapeutic vaccination (n=5). (C) Determination of stomach SIgA antibody against *H. pylori* lysates after therapeutic vaccination (n=5). (D) Quantification of gastric *H. pylori* colonization by CFU counting. Data are mean ± S.D. ***p < 0.001, **p < 0.01, ns, not significant.

The relevant cytokines IFN-γ, IL-4, and IL-17 in the supernatants of splenic lymphocyte cultures were determined using ELISA. *H. pylori* lysates significantly induced higher levels of IFN-γ (Figure 6 A), IL-4 (Figure 6 B), and IL-17 (Figure 6 C) in splenic lymphocytes from mice immunized with CTB-S3 or CTB-S5 than those from mice immunized with CTB. There were no significant differences at the levels of IFN-γ, IL-4, and IL-17 cytokines between CTB-S3 and CTB-S5 groups.

**FIGURE 6.**
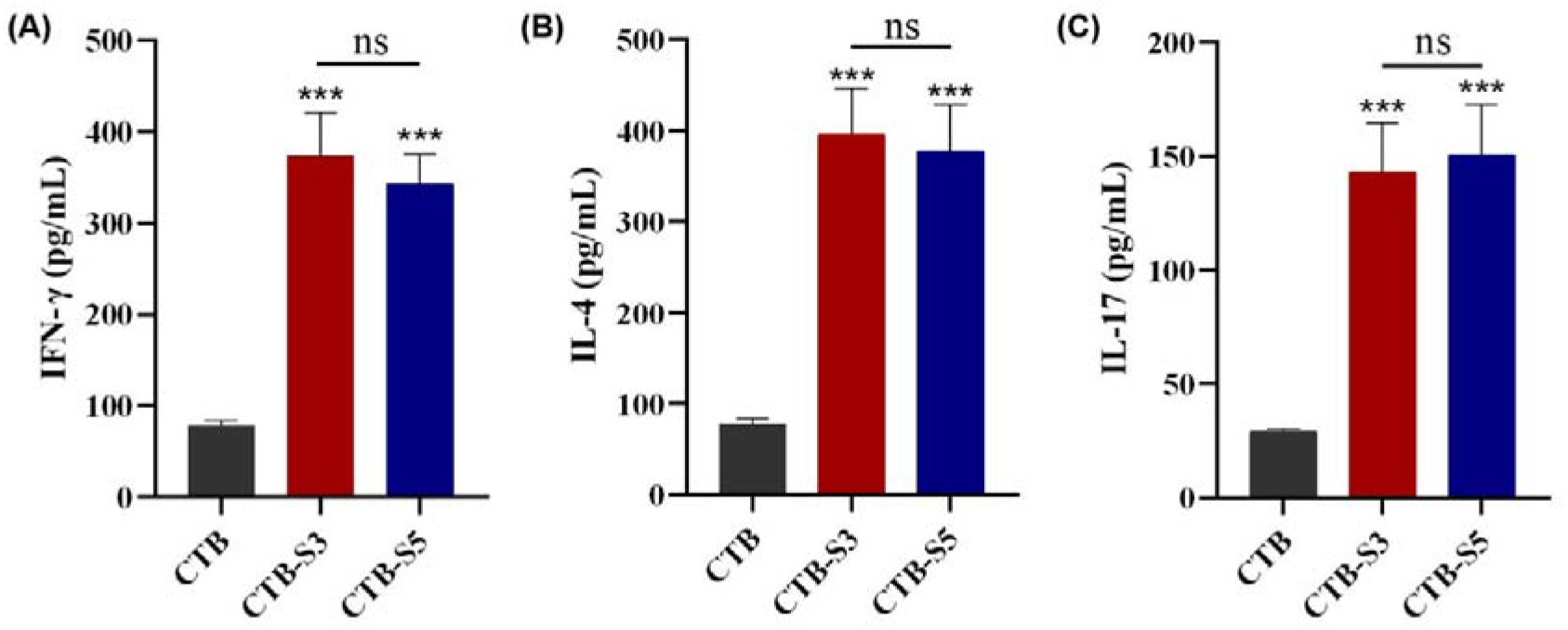
Cytokine production of splenic lymphocytes from *H. pylori*-infected mice after therapeutic immunization with CTB, CTB-S3, or CTB-S5. The cytokines including IFN-γ (A), IL-4 (B), and IL-17 (C) were detected by ELISA. Data are mean ± S.D. ***p < 0.001, ns, not significant.

Histology of the gastric biopsies revealed a severe glandular stomach ulcer and the inflammatory cell infiltration in the stomachs from the mice immunized with CTB. However, a mild or moderate degree of inflammatory cell infiltration was found in the stomachs from mice immunized with CTB-S3 or CTB-S5. Typical histological images of the stomach from different groups were shown in Figure 7. The histological results showed that the immunization with CTB-S3 or CTB-S5 could protect the *H. pylori*-infected stomach to a certain extent.

**FIGURE 7.**
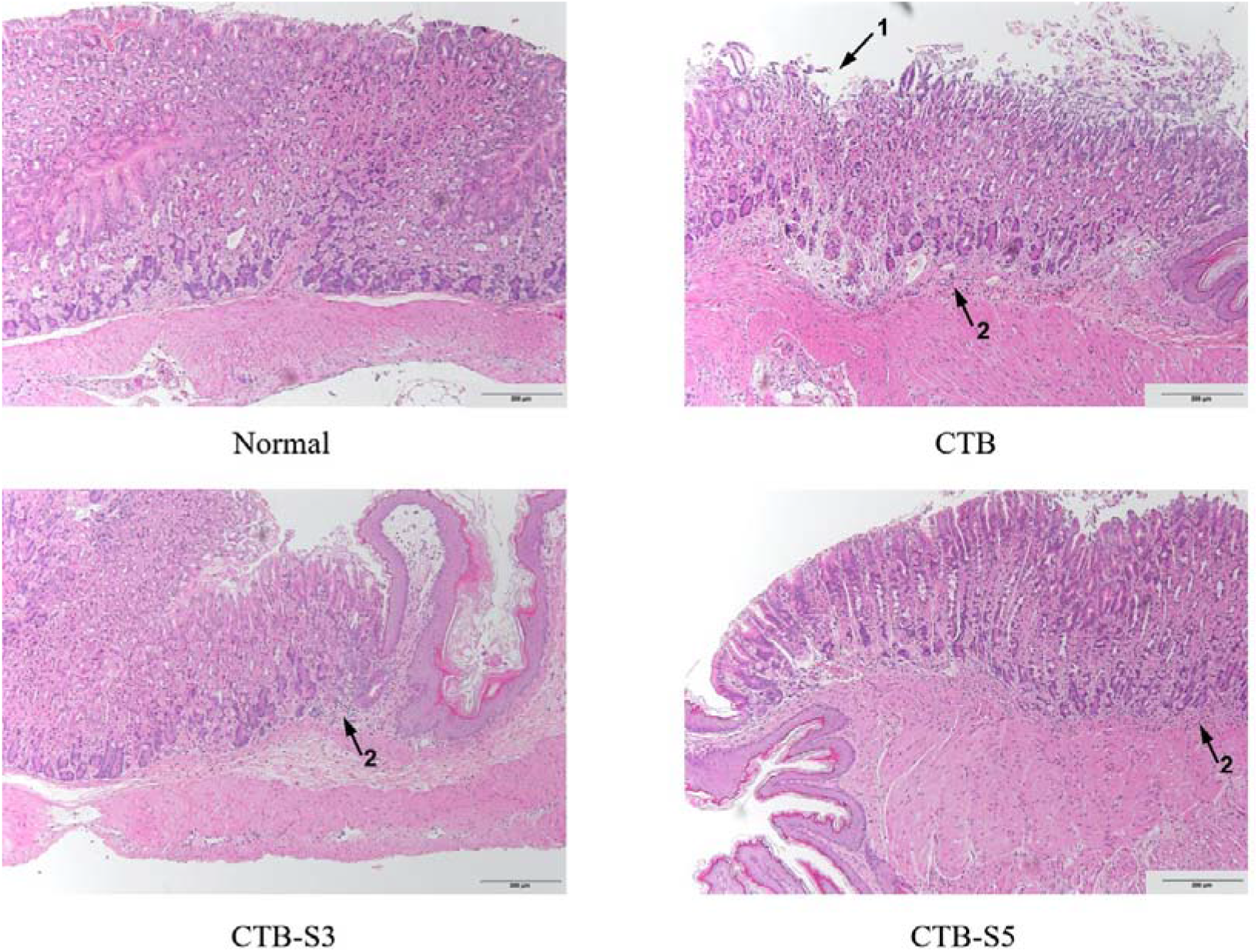
Gastric histology after therapeutic vaccination with CTB, CTB-S3, or CTB-S5. 1, Glandular stomach ulcer; 2, Inflammatory cell infiltration.

## 4| DISCUSSION

Since *H. pylori* resistance to antibiotic regimens is increasing, vaccination is becoming an increasingly important alternative therapy to control *H. pylori* infection. To solve the problem of the limited immune response induced by a single antigen, we developed a multi-epitope vaccine based on the epitopes from 5 *H. pylori* antigens including UreB, FlaA, AlpB, SabA, and HpaA. The epitopes with different permutations produced 7 different constructs after the primary screening of VaxiJen. Although identical in amino acid composition, their structures showed significant differences in structural folding and reliability (Figure S1 B; Table 2). The reliability and rationality of the predicted structure determined the stability of the recombinant multi-epitope antigen. Finally, Construct S3 and S5 performed well in structure validation were screened. At the beginning of our experiment, the expressions of tandemly linked constructs were failed; the successful expressions of CTB-S3 and CTB-S5 preliminarily confirmed the rationality of the multi-epitope vaccine designed by computer aid (Figure 3 B).

The construction of a multi-epitope vaccine aimed at inducing a broadly reactive antibody response to block multiple pathogenic channels of pathogens^36^. In our design, we chose chemotaxis-related antigens including UreB, FlaA, and adhesin-related antigens including AlpB, SabA, HpaA. In theory, the epitopes from the above antigens could induce the specific antibody response to inhibit *H. pylori* chemotaxis, adhesion, and transmission. After subcutaneous immunization, CTB-S3 and CTB-S5 vaccines could generate an extensive antibody response against 5 antigens including UreB, FlaA, AlpB, SabA, and HpaA (Figure 4 A). Thus, the epitope-specific antibodies may play a role in controlling *H. pylori* infection. Further, the purified anti-CTB-S3 IgG and anti-CTB-S5 IgG could inhibit the enzyme activity of *H. pylori* urease and *H. pylori* adhesins to AGS cells (Figure 4 C, D). These neutralizing antibodies are essential for inhibiting *H. pylori* colonization and pathogenicity. Besides, the oral immunization with CTB-S3 or CTB-S5 could induce a high level of *H. pylori*-specific serum IgG antibody (Figure 5 A), indicating that the epitope-specific antibody induced by oral immunization effectively targeted *H. pylori*.

To control the gastrointestinal pathogen infection, the high level of SIgA antibody induced by mucosal immunity was firstly considered; so the *H. pylori* vaccines are usually immunized intragastrically or intranasally^37^. In addition, CTB could enhance mucosal immune efficiency as a safe and efficient mucosal adjuvant^18,38^. In our study, in the form of intramolecular adjuvant, CTB was fused with Construct S3 or S5 to achieve co-expression. The CTB-S3 or CTB-S5 proteins still could bind the GM1 (Figure 3 D), indicating that CTB in the fused construct could play the adjuvant effect of promoting antigen uptake. The fused CTB-S3 or CTB-S5 could induce a high level of *H. pylori*-specific SIgA antibody (Figure 5 B). It is thoughtless to perform the structural prediction of the fusion protein of CTB and the multi-epitope antigen in some recent studies^39,40^. Since only one parsed protein structure could be used as a template, the structure prediction platforms based on parsed structures were not suitable for predicting the structures of fusion proteins^24^.

The CD4^+^ T cell immune responses, especially Th1, Th2, and Th17-biased responses are crucial for controlling *H. pylori* infection^41^. In our design, 3 T cell epitopes from UreB (UreB_409-421_, UreB_237-251_, and UreB_548-561_) were added into Construct CTB-S3 or CTB-S5. These 3 T cell epitopes were proved to be effective in the previous studies^11,12^. *H. pylori* lysates significantly induced higher levels of IFN-γ, IL-4, and IL-17 in splenic lymphocytes from mice immunized with CTB-S3 or CTB-S5 (Figure 6), indicating that the oral immunization with CTB-S3 or CTB-S5 could induce *H. pylori*-specific CD4^+^ T cell immune response.

In conclusion, we developed 2 multi-epitope vaccines, named CTB-S3 and CTB-S5 based on structural validation. The immunization with CTB-S3 or CTB-S5 could induce broad antigen-specific antibodies and neutralizing antibodies against *H. pylori* urease and adhesins. The oral immunization with CTB-S3 or CTB-S5 could significantly reduce *H. pylori* colonization and protect the stomach in the *H. pylori*-infected mice, which may be related to mixed epitope-specific antibody responses and Th cell immune response. The CTB-S3 and CTB-S5 vaccine may be promising vaccine candidates to control *H. pylori*-infection and the vaccine design strategy based on computer aid will provide a reference for the vaccine construction of other pathogens.

## ACKNOWLEDGEMENTS

This work was supported by Science and Technology innovation Plan of Shanghai (19391902000). The funders did not participate in study design, data collection, analysis, and interpretation or preparation of the manuscript.

## AUTHOR CONTRIBUTIONS

Study concept and design: Junfei Ma and Qing Liu. Analysis and interpretation of data: Junfei Ma, Shuying Wang, and Qianyu Ji. Drafting of the manuscript: Junfei Ma and Shuying Wang. Study supervision: Qing Liu.

## DATA AVAILABILITY STATEMENT

All data generated or analysed during this study are included in this published article (and its supplementary information files).

## SUPPORTING INFORMATION

**FIGURE S1.**
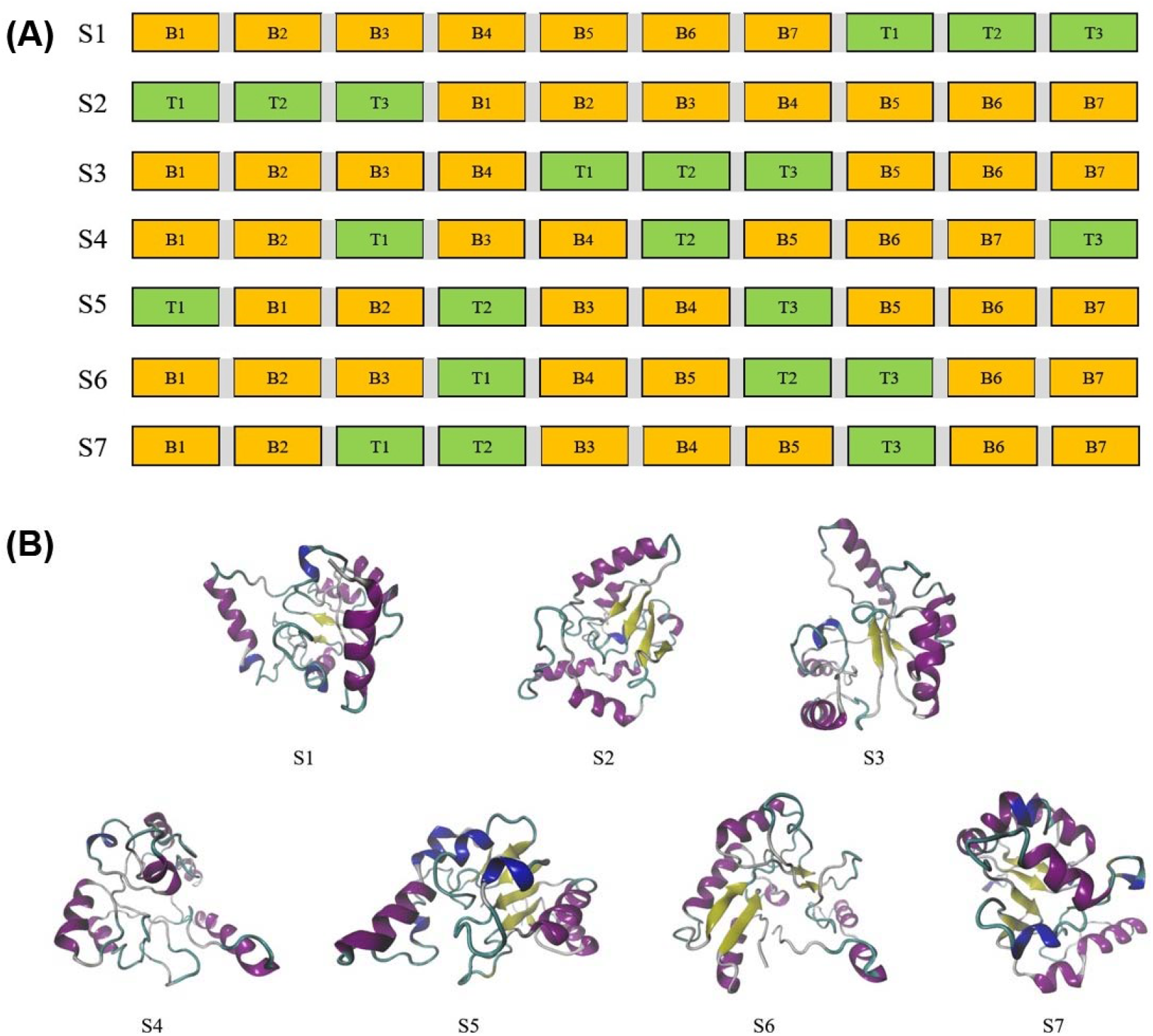
The design of multi-epitope antigens. (A) The constructs of multiepitope antigens (Numbered S1-S7) with 7 random permutations. B cell epitopes filled with Color Orange and T cell epitopes filled with Color Green. (B) The structures of the 7 constructs predicted by I-TASSER.

**TABLE S1.**
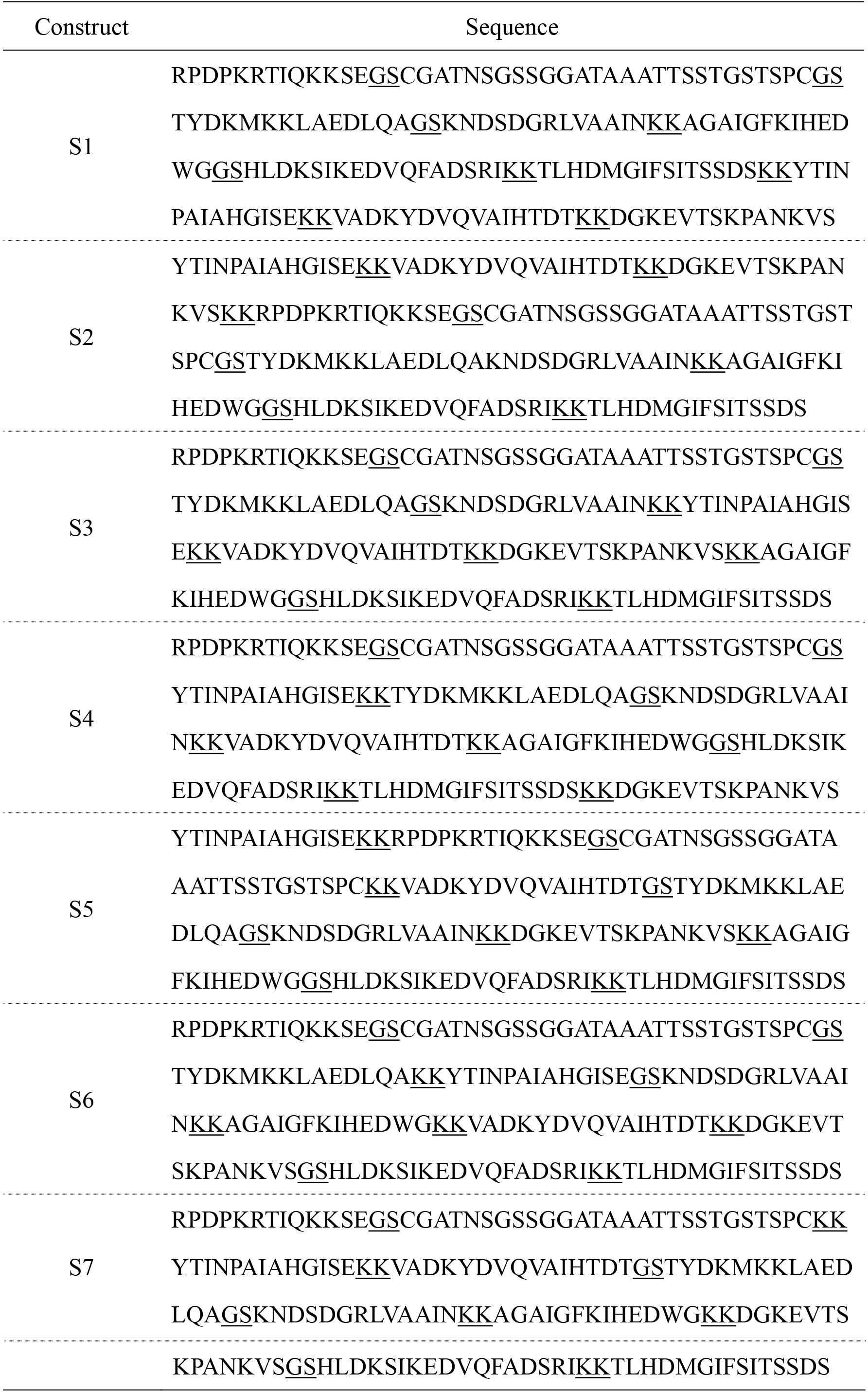
The sequence of 7 constructs S1-S7.

